# Evaluating the potential benefits and pitfalls of combining protein and expression quantitative trait loci in evidencing drug targets

**DOI:** 10.1101/2022.03.15.484248

**Authors:** Jamie W Robinson, Thomas Battram, Denis A Baird, Philip C Haycock, Jie Zheng, Gibran Hemani, Chia-Yen Chen, Tom R Gaunt

**Author notes:** Corresponding author: Jamie Robinson –. Oakfield House, Oakfield Grove, Clifton, Bristol, BS8 2BN.

## Abstract

Molecular quantitative trait loci (molQTL), which can provide functional evidence on the mechanisms underlying phenotype-genotype associations, are increasingly used in drug target validation and safety assessment. In particular, protein abundance QTLs (pQTLs) and gene expression QTLs (eQTLs) are the most commonly used for this purpose. However, questions remain on how to best consolidate results from pQTLs and eQTLs for target validation.

In this study, we combined blood cell-derived eQTLs and plasma-derived pQTLs to form QTL pairs representing each gene and its product. We performed a series of enrichment analyses to identify features of QTL pairs that provide consistent evidence for drug targets based on the concordance of the direction of effect of the pQTL and eQTL. We repeated these analyses using eQŢLs derived in 49 tissues.

We found that 25-30% of blood-cell derived QTL pairs have discordant effects. The difference in tissues of origin for molecular markers contributes to, but is not likely a major source of, this observed discordance. Finally, druggable genes were as likely to have discordant QTL pairs as concordant.

Our analyses suggest combining and consolidating evidence from pQTLs and eQTLs for drug target validation is crucial and should be done whenever possible, as many potential drug targets show discordance between the two molecular phenotypes that could be misleading if only one is considered. We also encourage investigating QTL tissue-specificity in target validation applications to help identify reasons for discordance and emphasise that concordance and discordance of QTL pairs across tissues are both informative in target validation.

## Main Text

Millions of genetic variants have now been identified as being associated with a variety of diseases. The protein product of proximal genes to these genetic variants are more likely to be successful drug targets than proteins without proximal genetic variation related to the disease of interest ^1^. It follows that if the supporting disease-associated genetic variants are also associated directly with protein abundance, the protein will be more likely to be a successful drug target ^1–3^. However, a genetic variant may have different effects on the expression level of a protein coding gene and the actual protein abundance. Variants with discordant genetic effects on gene expression levels and protein abundance are difficult to interpret and are frequently de-prioritised in target validation. So-called “opposite eQTL effects” have been previously observed in tissue-specific gene regulation ^4^. however, few studies have investigated concordance of QTLs across different omic layers and how this may affect drug target validation and prioritisation. Using protein quantitative trait loci (pQTLs) from the recently published Fenland study ^5^ and Zhang, et al ^6^, and expression quantitative trait loci (eQŢLs) from eQTLGen ^7^ and expanded tissue data from GTEx v8 ^8^, we investigated whether discordance of QŢL effects influenced the validity of proteins as drug targets and what might be responsible for this discordance. We proposed two hypotheses to guide this work: firstly, for a given gene, if the pQTL and eQŢL are concordant, the protein encoded by the gene will be more likely to be a valid drug target; secondly, the discordance between pQTL and eQŢL effects can be partially explained by tissue differences in which these data are measured. For example, many blood pQTLs are based on the abundance of proteins which are commonly secreted to the blood from other tissues (i.e. these proteins form part of the secretome ^9^) whereas blood eQTLs are generally derived from blood cells^7;8^.

We extracted 3,764 conditionally independent cis-pQTLs (within 1Mb of the gene coding region) derived in plasma for 1,693 proteins from their original publications (Fenland study: 3,498 pQTLs for 1,627 proteins; Zhang, *et al*: 266 non-overlapping pQTLs for 66 proteins)^5;6^. We constructed QTL pairs by performing a one-to-one SNP lookup for eQTLs from eQTLGen ^7^, allowing for proxy SNPs (r^2^ > 0.8) to increase power, and harmonising beta estimates to the same effect allele. SNPs were subjected to a P value threshold for selection adjusted for the number of available proteins (P < 2.95×10^-5^,0.05/1693). This threshold was applied to both the pQTL and eQTL P value for each QTL pair.

Generally, many of the proteins had at least one associated pQTL. To avoid sampling bias due to proteins with more than one associated pQTL, we categorised the QTL pairs as either primary or non-primary. Primary QTL pairs were defined as those QTL pairs where the pQTL had the lowest P value and non-primary QTL pairs were defined as all other QTL pairs. Primary status was determined before creating QTL pairs, so it was possible for some proteins to have only non-primary QTL pairs if the primary pQTL was not paired with an eQTL. We made use of only the primary QTL pairs in the main analyses, and we conducted sensitivity analyses with both primary and non-primary QTL pairs. The pQTL studies used SomaLogic SOMAmers, which can suffer bias in measurement due to differential binding to non-synonymous variants in the target sequence ^5;10^. To address this, we dropped QTL pairs which included an indel pQTL as it is expected that indels have a stronger effect on protein structure, and hence binding affinity, compared to SNPs.

To determine concordance between QTL pairs, we conducted a naïve sign test comparing the beta estimates between pQTLs and eQTLs **(Figure 1).** However, we were concerned that discrepancies in statistical power might lead to observing discordance between QTL pairs so we utilised a Bayesian winner’s curse correction (described previously in Okbay, *et al.* ^11^; code supplied as supplement) to check if this was the case. Results from the winner’s curse corrected analysis determined whether differences in the signs of the beta estimates was due to lack of power and were used to determine whether the definition of concordance from the naïve test was biased.

**Figure 1.**
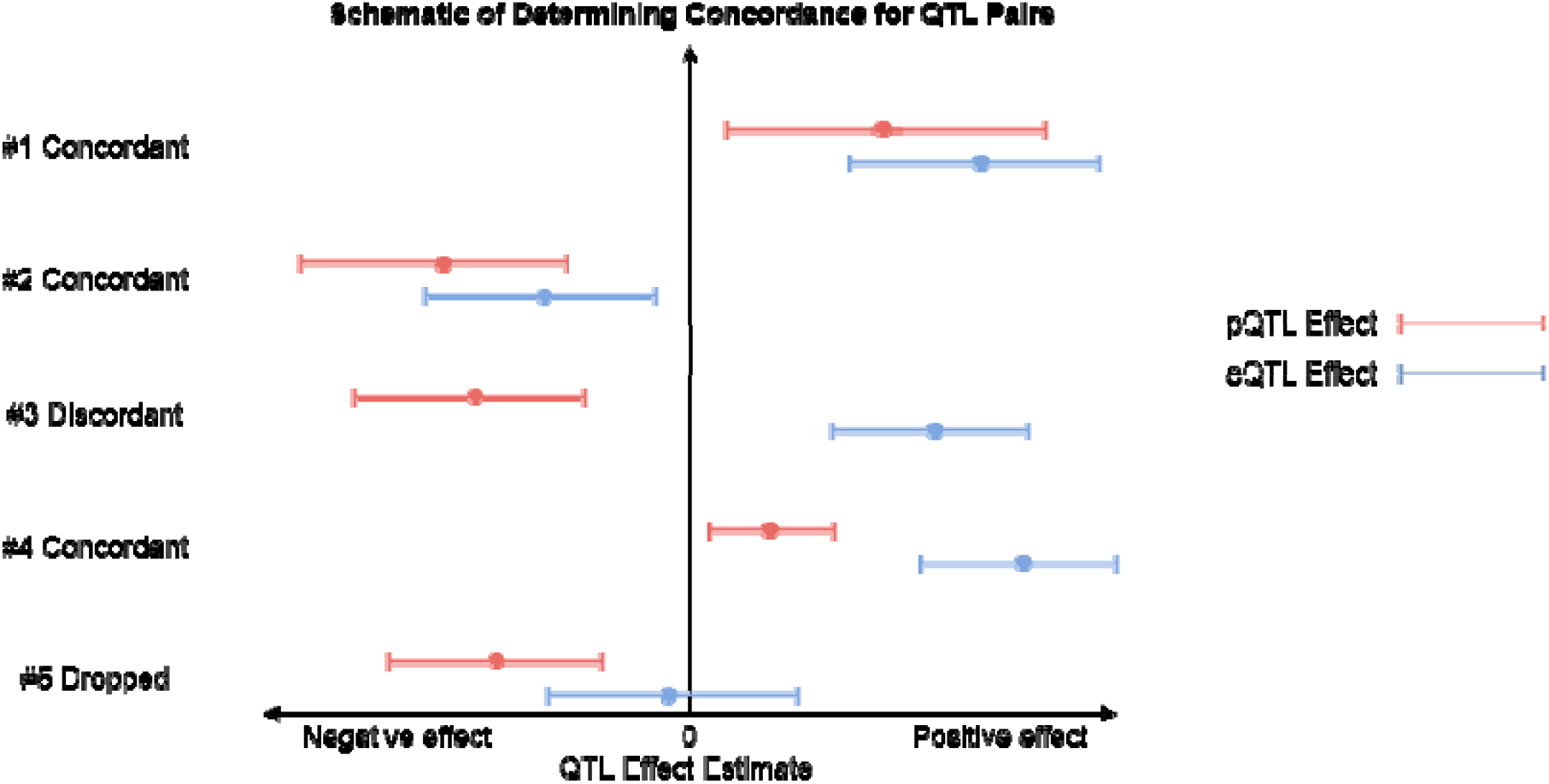
Schematic showing how concordance between pQTL and eQTL pairs was determined using a naïve sign test. QTL pairs whose direction of effect for the beta estimates were in agreement were classified as concordant, regardless of whether that effect was positive or negative (cases #1 and #2). In case #3, where the direction of effects do not agree then this QTL pair was classified as discordant. It is important to note that our definition of concordance did not exclude nonoverlapping effect estimates for the two QTLs, so long as their beta estimates were in the same direction of effect (case #4). Finally, for case #5, where one of the QTLs is consistent with the null hypothesis according to our threshold (P < 2.95×10^-5^, 0.05/1,693), these were dropped from the analysis.

In the enrichment analysis, we utilised the categorisation of concordance for QTL pairs to determine whether terms from the Drug-Gene Interaction database (DGIdb) ^12^ are enriched for concordant or discordant QTL pairs. Terms included were those that are drug target relevant such as the “druggable genome”, that annotates genes which encode druggable proteins and are likely to be clinically relevant^2;13–15^. P values for this analysis were calculated using Fisher’s exact test on a 2×2 contingency table for each term present in the enrichment analyses (concordant/discordant against linked to term/not linked to term). We present both unadjusted and false discovery rate (FDR)-adjusted P values based on the number of terms present in DGIdb. However, given the hierarchical structure and overlap between the DGIdb terms, FDR-adjusted P values are likely to be conservative.

Finally, as we hypothesised that discordance between QTLs may arise due to differences in the tissue where the QTLs are measured, we constructed QTL pairs using eQTLs across 49 tissues present in GTEx v8 ^8^. The construction strategy was similar to the main analysis, except that we did not search for proxy SNPs in the GTEx dataset as there was comparable coverage between QTL pairs using eQTLs from blood and other tissue eQTLs; however, there were fewer QTL pairs available at the selection threshold of P < 2.95×10-^5^ due to lower power which varied by tissue (**Table S1**). We repeated both concordance tests for the GTEx QTL pairs.

A potential reason for discordance in QTL pairs constructed with GTEx eQTLs could be that plasma proteins are generally secreted to blood from the tissue where the gene is primarily expressed. Therefore, it could be expected that the abundance of a plasma protein would be more correlated to the expression of its coding gene within the tissue that secretes that protein into the plasma, rather than its expression in other tissues. While many plasma proteins originate from the liver, they can potentially be secreted from many other tissues ^9;10^. To ascertain whether proteins without evidence of secretion from specific tissues to plasma (instead of, for example, to the intra-cellular region) were driving discordance, we combined GTEx QTL pairs with secretome evidence from the Human Protein Atlas (HPA)^9^. Specifically, QTL pairs were linked to those proteins with evidence of being secreted to plasma from a tissue which matched with a GTEx eQTL that formed the QTL pair.

There were a total of 1,296 QTL pairs for 838 proteins constructed using plasma-derived pQTLs from the Zhang, *et al*. and Fenland studies and blood-cell derived eQTLs from eQTLGen which met the P value threshold of P < 2.95×10^-5^ (both the pQTL and eQTL had to pass this threshold) (**Tables S1** and S**2**). Of these, 736 QTL pairs were classified as primary and the remaining 560 QTL pairs as non-primary (not all 838 proteins had a primary QTL pair because the primary pQTL was not paired with eQTL). We observed a concordance rate of 71.8% for primary QTL pairs and 72.4% for all QTL pairs using the naïve concordance test **(Figure 1).** In the Bayesian winner’s curse analysis with primary QTL pairs only, all 736 primary QTL pairs were expected to be significant at the 5% level and to have matching signs. However, only 528 QTL pairs (71.8%) were observed to have a matching sign although all 736 QTL pairs were observed to be significant at the 5% level. A similar pattern of concordance rate after correction for winner’s curse was seen for the all QTL pairs analysis **(Table S3).** Therefore, the results from the Bayesian winner’s curse correction analysis agreed with the naïve concordance test which indicated that concordance was not affected by differential power between the pQTL and eQTL datasets.

We performed a *post hoc* analysis to determine if the rate of concordance differed for primary and non-primary signals. There was no significant difference in the pattern of concordance between the primary and non-primary QTL pairs (primary QTL pairs: concordant = 528, discordant = 208; nonprimary QTL pairs: concordant = 410, discordant = 150; Fisher’s exact test P = 0.57).

For the enrichment analysis, we observed that primary discordant QTL pairs were enriched for the terms “cell surface” (P = 0.003, P_FDR_ = 0.07) and “protease” (P = 0.014, P_FDR_ = 0.15) (*Figure 2i*). Including non-primary QTL pairs showed enrichment for discordant QTL pairs with the term “protein phosphatase” (P = 0.022, P_FDR_ = 0.48) (*Figure 2ii*). The term “druggable genome”, which is an umbrella term for many of the terms DGIdb collate, was not enriched for either concordant or discordant QTL pairs. There was no longer any evidence for enrichment for the terms “cell surface” and “protease”.

**Figure 2.**
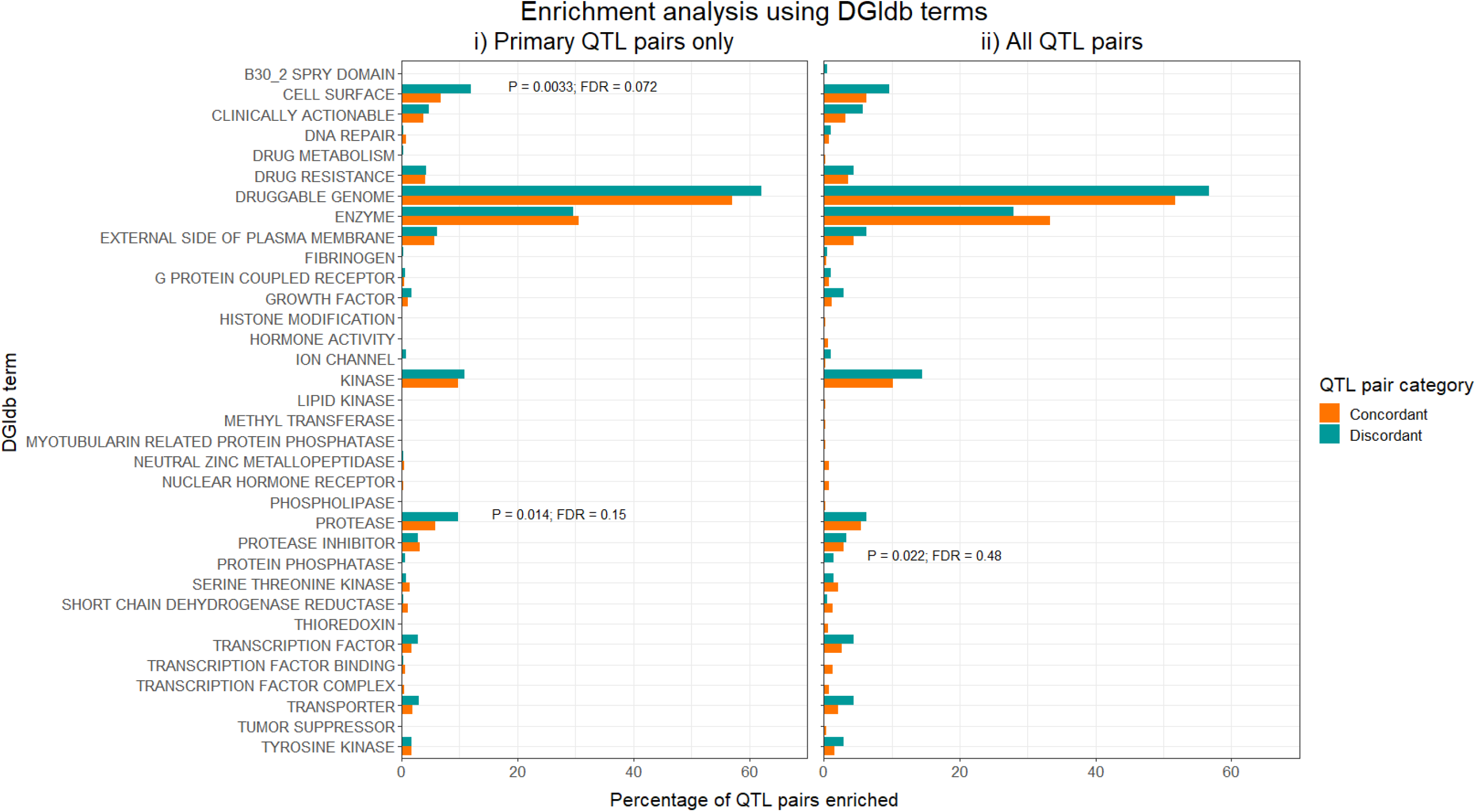
Results for the enrichment analysis using the Drug-Gene Interaction database (DGIdb) for druggability-related terms. The bars show the percentage of concordant or discordant QTL pairs which were enriched for a given term. P values were calculated using Fisher’s exact test, an d unadjusted and FDR-adjusted P values are shown for those terms which reached at least nominal significance.

We further hypothesised that discordance between QTL pairs may arise due to tissue differences in the datasets used to estimate the effects of gene expression and protein abundance. To examine this, we combined the same plasma pQTLs with GTEx eQTLs, which we refer to as GTEx QTL pairs to differentiate from the blood QTL pairs **(Table S4).** There were 12,923 GTEx QTL pairs after selection (pQTL and eQTL P value < 2.95×10^-5^), of which 8,730 were classified as primary and 4,193 were non-primary. Concordance rates varied across individual tissues, where primary QTL pairs consisting of eQTLs measured in the kidney (100.0%), uterus (96.8%) and small intestine (93.5%) showed the highest concordance rate, while the cerebellum (69.0%), cerebellar hemisphere (74.3%) and testis (76.4%) showed the lowest rate of concordance **(Figures 3i** and **3ii**). Overall, the mean concordance rate for primary GTEx QTL pairs was 85.6% and for all GTEx QTL pairs was 83.6%. Combining GTEx QTL pairs with HPA secretome evidence, there were 111 QTL pairs with such evidence and which met the P value threshold of 2.95×10^-5^. These 111 QTL pairs consisted of 66 primary and 45 non-primary pairs across 23 tissues. Using the subset of GTEx QTL pairs which had secretome evidence, we observed a concordance rate of 90.9% for primary pairs and 86.5% for all pairs. There were too few pairs to robustly examine concordance on a per-tissue level **(Figure 4).**

**Figure 3.**
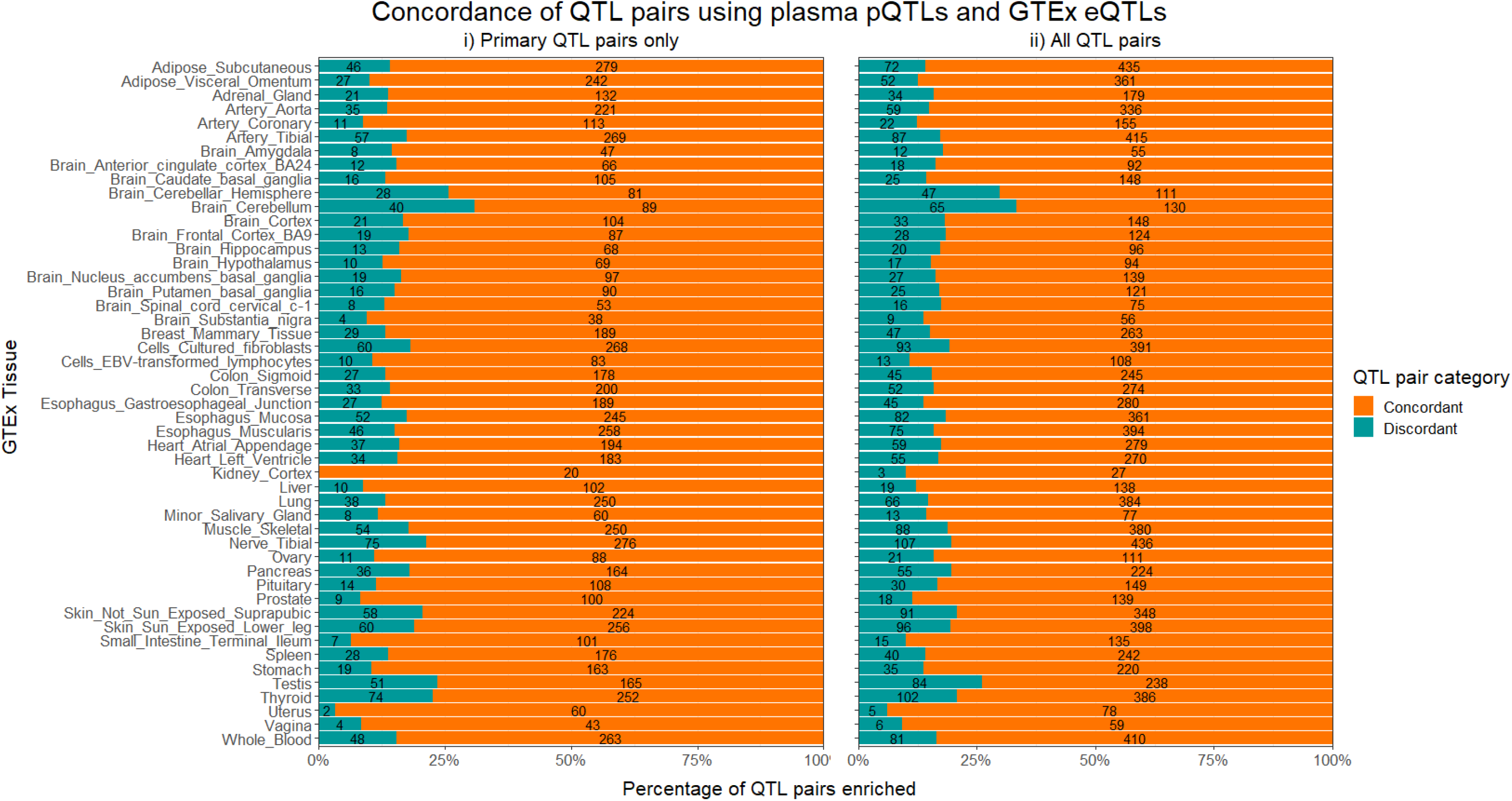
Concordance rates for QTL pairs formed of plasma pQTLs and GTEx eQTLs. Number of concordant or discordant QTL pairs are given as numbers in the bars.

**Figure 4.**
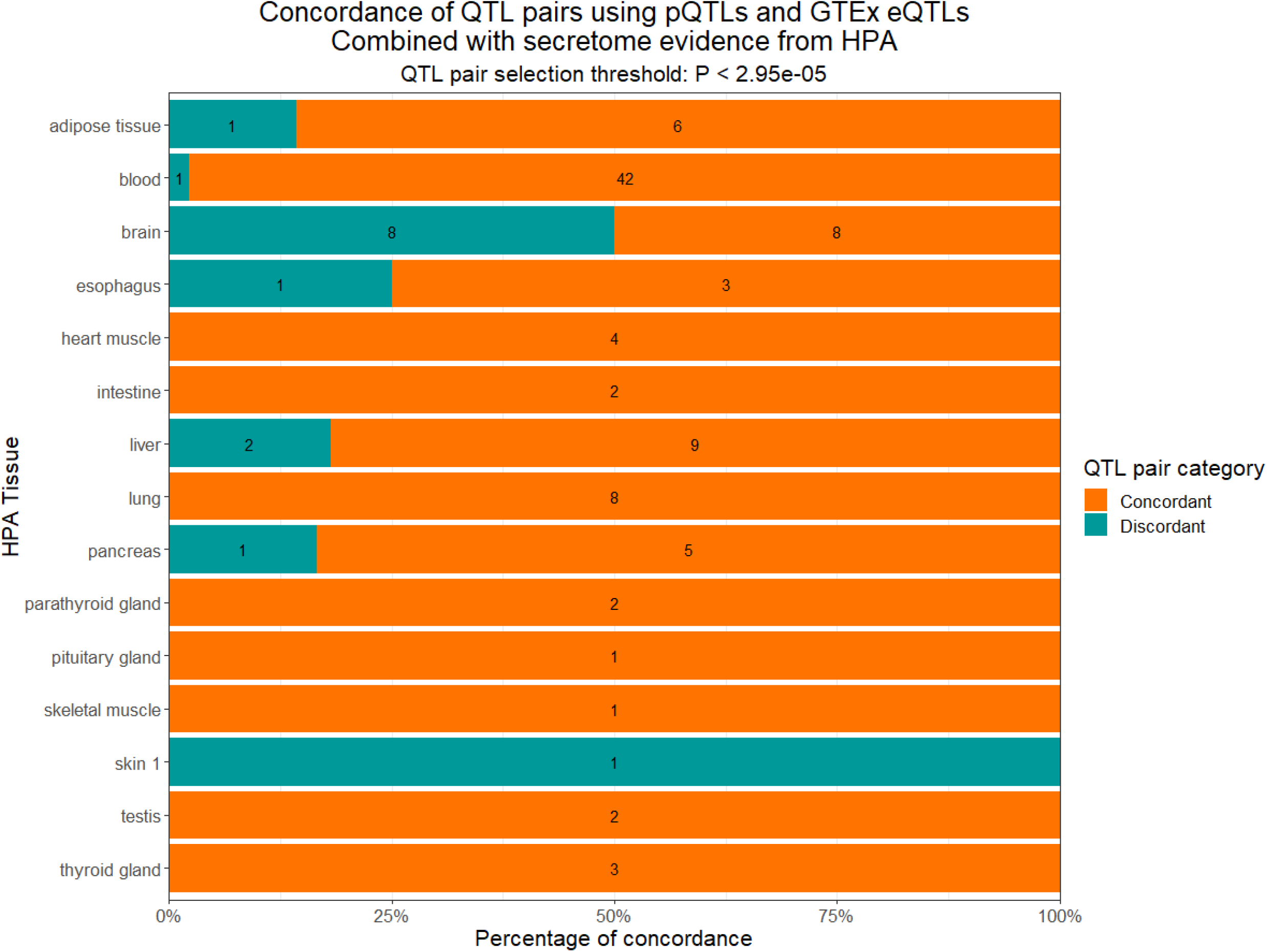
Concordance rates for all QTL pairs (primary and non-primary) with HPA secretome evidence (i.e. the protein is secreted to plasma). These data consisted of plasma-derived pQTLs and GTEx eQTLs and were combined with secretome evidence from HPA. Number of QTL pairs are given as numbers in the bars.

As concordance rates across tissues were broadly similar, we conducted a *post hoc* analysis to determine if this was because the GTEx eQTLs showed similar effects across tissues. We performed a test for heterogeneity between the GTEx eQTLs for each tissue and found strong evidence for homogeneity of eQTL effect estimates **(STable 5).**

We proposed two hypotheses to guide our work: firstly, proteins encoded by genes with a concordant pQTL and eQTL would more likely be druggable; and secondly, tissue differences may partially explain discordance between pQTL and eQTL effects for corresponding gene products.

Initially, we paired plasma-derived pQTLs with blood cell-derived eQTLs and observed a concordance rate of 71.7% for primary QTL pairs which reached a threshold of P < 2.95×10^-5^. As there was lower statistical power to identify pQTLs than eQTLs, we implemented a sensitivity analysis to determine whether discordance between QTLs were due to this difference in power. We found that all QTL pairs’ P values were expected and observed to be significant at the 5% level but that the same percentage of QTL pairs were discordant as in the naïve concordance test. Therefore, discordance between the plasma-derived pQTLs and blood cell-derived eQTLs was not due to the differences in sample sizes from which these data were measured. We noted that concordance rates between GTEx QTL pairs ranged from 69.0% to 100.0% and the mean overall concordance rate for all GTEx QTL pairs was 83.6%. Concordance between GTEx QTL pairs increased further to 90.9% when combined with secretome evidence (though there were only 111 pairs included in this comparison).

A previous study conducted a similar concordance test for eQTLs in GTEx and found that tissue-dependent discordant eQTLs effects were present for 2,323 genes of the 31,212 they analysed (7.4%)^4^. Our study is the first to have looked at concordance rates between different omics layers. Whilst our analyses included a relatively smaller number of proteins/genes, we observed that concordance rates between pQTLs and eQTLs increased as tissue differences were taken into account, e.g. by using tissue-derived eQTLs and combining QTL pairs with secretome evidence which better captures the effects of plasma-derived pQTLs. However, there appeared to be a consistent set of QTL pairs which did not show concordance between pQTL and eQTL effects. By way of example, the *HP* gene (which encodes haptoglobin) is well-characterised and has been shown to be highly expressed in the liver and secreted to blood where its primary function of binding free haemoglobin takes place ^16–18^. However, we found that the genetic effect on *HP* expression measured in liver tissue was discordant with the genetic effect on protein abundance measured in blood plasma.

We conceived of the following possible explanations for why QTL pairs might be discordant, even when considering tissue differences in the QTLs: 1) measurement error; for example, a variant may positively affect gene expression but negatively affect SOMAmer binding efficacy; 2) the QTL effect is simultaneously affecting transcript stability and translation efficiency in opposite directions; for example, a variant may reduce translational efficiency and could lead to a “backlog” of transcripts; 3) temporal effects; for example, the protein product of a gene may be more persistent or stable whilst the expression of the gene oscillates. More broadly, point 2 is an example of how molecular pleiotropy can manifest and cause discordance across different omics layers. Considering the example of the *HP* gene, although the biology surrounding *HP* is well-studied and the path from gene to protein is clear, the effects of genetic variants on these biological processes might still differ. The functional characterisation of molecular QTLs is important to understanding discordance between types of QTLs.

We also hypothesised that those proteins whose QTLs were concordant would have greater evidence of being druggable; however, our results suggested this was not the case. We observed no strong evidence for a difference in enrichment for the DGIdb “druggable genome” term between concordant and discordant QTLs. We did observe that discordant QTL pairs were enriched for “cell surface”, a Gene Ontology (GO) term to annotate proteins which attach to the plasma membrane or cell wall, “protease”, an enzyme which catalyses proteolysis which is likely to be the mechanism of effect of a druggable protein, and “protease inhibitor”, a class of drugs which target proteases; however, these generally contained small number of QTL pairs and should be treated with caution. It should also be noted that although the FDR-adjusted P value showed weak evidence, many of the DGIdb terms are correlated and so this adjustment is likely to be conservative. Given that both concordant and discordant QTL pairs are equally likely to be druggable, we advise that authors in future studies should not base their evidence on a single molecular phenotype, and should consider how to interpret the evidence from discordant QTL pairs as these may still be informative for evidencing drug targets.

The data we leveraged to perform these analyses were derived from large sample sizes; however, it is clear that the lack of tissue-specific pQTL data limited our ability to further test concordance across omics layers at a per-tissue level. Our selection criteria for QTLs meant that we minimised common sources of bias: we excluded trans-QTLs as these tend to be more pleiotropic than variants measured in the *cis* region ^3^. QTL pairs were categorised as primary or non-primary to address sampling bias; and pQTLs were independent as reported by the original authors (for example, as determined by conditional analyses).

However, there were some limitations to our study. Firstly, it is possible that discordance between QTL pairs could be due to distinct causal variants for gene expression and protein abundance that are in high LD with each other. Full summary statistics for the pQTL data were not available and so we could not conduct colocalisation analyses between the pQTL and eQTL data to test this. Secondly, there is the possibility of collider bias in the enrichment analysis **(Figure 5)**. This bias can be induced due to the selection criteria for a protein to be included in a proteomic panel, for example, SomaLogic’s SomaScan assay can currently measure up to 7,000 proteins, a small percentage of the overall human proteome. However, these proteins are likely to have been selected on particular characteristics potentially including, but not limited to, 1) uniqueness, where paralogues of proteins are harder to specifically measure; 2) sequence variation, i.e. it is harder to measure proteins with higher sequence variability; and 3) circulating levels, where proteins with low abundance or high temporal variability will be inherently harder to measure. Furthermore, it is likely that proteins, whose associated genes form part of the druggable genome, are more likely to have a protein assay exist to measure that protein as they will be informative for drug discovery efforts. Though unlikely, this can lead to an induced association which may bias the results from our enrichment analysis.

**Figure 5.**
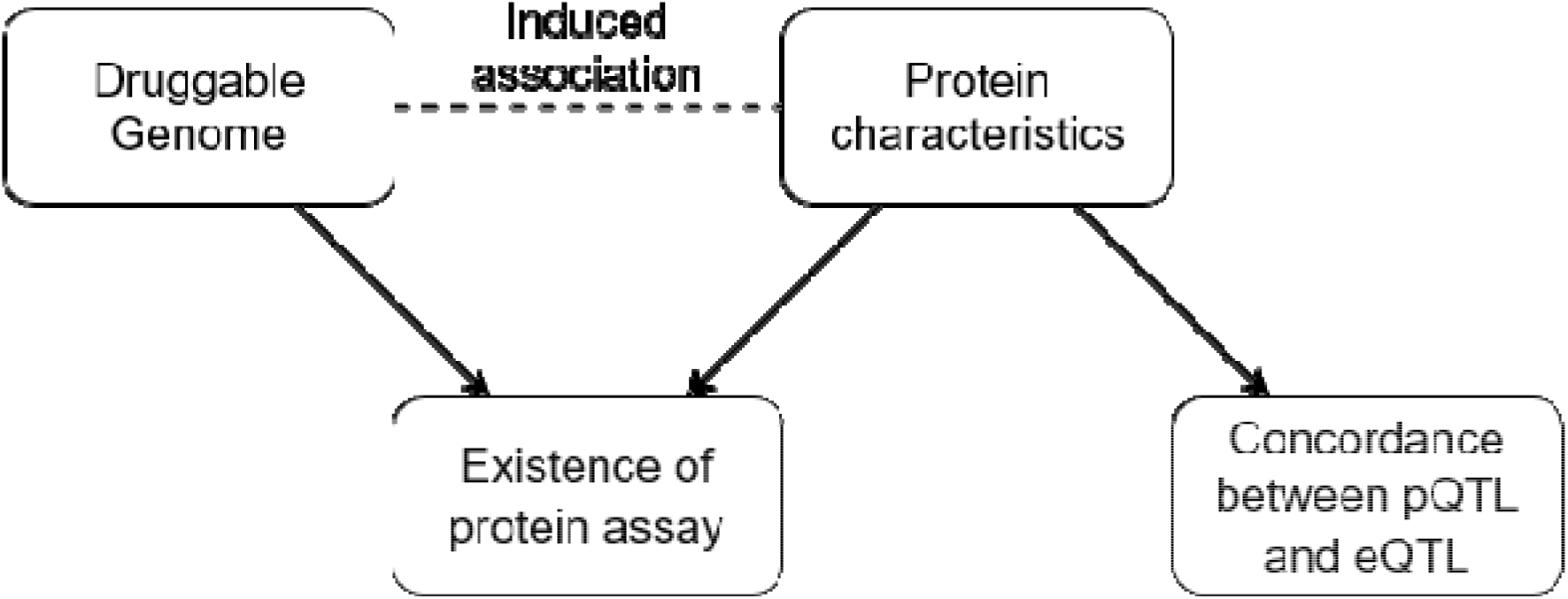
Directed acyclic graph showing how collider bias may be induced due to the existence of protein characteristics associated with both the existence of an assay to measure abundance of a protein and concordance between the pQTL and eQTL. Some examples of protein characteristics that may induce collider bias are given in the main text. For example, paralogues may influence the existence of an assay which measures the abundance of a protein and may also influence concordance between a pQTL associated with that protein and an eQTL associated with the encoding gene.

Our analyses showed that discordance between plasma-derived pQTLs and blood cell-derived eQTLs is common but can be mitigated when considering tissue differences in which these QTLs are measured. Utilising tissue-derived QTLs and considering the biological differences in how these QTLs are measured greatly increases concordance rates, implying these are relevant considerations to make in studies which integrate multiple omics layers. Importantly, our results suggested that discordant QTLs pairs are common for druggable genes and that a careful consideration of crosstissue effects is warranted to avoid unnecessarily discarding evidence for potential drug targets. As larger and more comprehensive pQTL datasets become available, so too will our ability to assess discordance between the genetic effects on gene expression and protein abundance. Integrating these data into multidimensional studies for evidencing drug targets will aid such efforts and lead to translational benefits.

## Data and Code Availability

All data used for this work are publicly available. pQTL data are available from the original publications (Fenland cohort: https://doi.org/10.1126/science.abj1541 and Zhang, *et al.*: https://doi.org/10.1101/2021.03.15.435533). eQTLGen and GTEx data are available on their websites https://www.eqtlgen.org/, and https://gtexportal.org/home/respectively. Our code used to generate these results is available at https://github.com/jwr-git/qtl_analysis.

## Supplemental Data

Six supplemental tables are provided in one Excel file. Supplemental note contains code to run the winner’s curse correction analysis.

## Acknowledgements

JWR and TRG receive funding from Biogen. JWR, TB and TRG receive funding from the UK Medical Research Council Integrative Epidemiology Unit at the University of Bristol (MC_UU_00011/4, MC_UU_00011/1). PCH is supported by CRUK Integrative Cancer Epidemiology Programme (C18281/A19169). GH is supported by the Wellcome Trust and Royal Society [208806/Z/17/Z], University of Bristol. The funders had no role in study design, data collection and analysis, decision to publish, or preparation of the manuscript.

