## Supplementary Code for "Evaluating the potential benefits and pitfalls of combining protein and expression quantitative trait loci in evidencing drug targets"

# Winner's curse replication analysis
# Originally described in https://doi.org/10.1038/nature17671
# Used to determine whether effects replicate across to GWAS at a given
# significance level.

#' Winner's curse replication analysis
#' 
#' @param b_disc Vector of beta estimates (discovery)
#' @param b_rep Vector of beta estimates (replication)
#' @param se_disc Vector of standard errors (discovery)
#' @param se_rep Vector of standard errors (replication)
#' @param alpha Alpha significance level, e.g. 0.05 for 95% confidence
#' 
#' @return list of results
#'   $res, tibble:
#'     Contains a 4x4 tibble containing columns for
#'     nsnp - number of SNPs
#'     metric - sign/p-value (which statistic is being tested)
#'     datum - expected/observed (according to significance level)
#'     value - number of SNPs which were expected/observed to replicate
#'  $variants, tibble:
#'     Contains a nsnpx8 tibble containing columns for:
#'     sig - significant
#'     sign - sign
#'     fdisc - F-statistic in discovery dataset
#'     frep - F-statistic in replication dataset
#'     p_rep - P value replication
#'     b_disc - Beta estimate in discovery dataset
#'     b_rep - Beta estimate in replication dataset
#'     same_sign - (logical) Does sign of beta estimates agree?
prop_overlap = function(b_disc, b_rep, se_disc, se_rep, alpha)
{
  p_sign <- pnorm(-abs(b_disc) / se_disc) * pnorm(-abs(b_disc) / se_rep) + ((1 - pnorm(-abs(b_disc) / se_disc)) * (1 - pnorm(-abs(b_disc) / se_rep)))
  p_sig <- pnorm(-abs(b_disc) / se_rep + qnorm(alpha / 2)) + (1 - pnorm(-abs(b_disc) / se_rep - qnorm(alpha / 2)))
  p_rep <- pnorm(abs(b_rep)/se_rep, lower.tail=FALSE)
  res <- dplyr::tibble(
    nsnp=length(b_disc),
    metric=c("Sign", "Sign", "P-value", "P-value"),
    datum=c("Expected", "Observed", "Expected", "Observed"),
    value=c(sum(p_sign, na.rm=TRUE), sum(sign(b_disc) == sign(b_rep)), sum(p_sig, na.rm=TRUE), sum(p_rep < alpha, na.rm=TRUE))
  )
  return(list(res=res, variants=dplyr::tibble(
    sig=p_sig, 
    sign=p_sign, 
    fdisc=b_disc^2/se_disc^2, 
    frep=b_rep^2/se_rep^2, 
    p_rep=p_rep, 
    b_disc,
    b_rep,
    same_sign=sign(b_disc) == sign(b_rep))))
}
